# Ancient and recent introgression shape the evolutionary history of pollinator adaptation and speciation in a model monkeyflower radiation (*Mimulus* section *Erythranthe*)

**DOI:** 10.1101/2020.09.08.287151

**Authors:** Thomas C. Nelson, Angela M. Stathos, Daniel D. Vanderpool, Findley R. Finseth, Yao-wu Yuan, Lila Fishman

**Author notes:** corresponding authors (TCN), (LF). Embark Veterinary, Inc., Boston, Massachusetts, United States of America (Thomas C. Nelson). Keck Science Department, Claremont-McKenna, Scripps, and Pitzer Colleges, Claremont, California, United States of America (Findley R. Finseth).

## Abstract

Inferences about past processes of adaptation and speciation require a gene-scale and genome-wide understanding of the evolutionary history of diverging taxa. In this study, we use genome-wide capture of nuclear gene sequences, plus skimming of organellar sequences, to investigate the phylogenomics of monkeyflowers in *Mimulus* section *Erythranthe* (27 accessions from seven species). Taxa within *Erythranthe*, particularly the parapatric and putatively sister species *M. lewisii* (bee-pollinated) and *M. cardinalis* (hummingbird-pollinated), have been a model system for investigating the ecological genetics of speciation and adaptation for over five decades. Across >8000 nuclear loci, multiple methods resolve a predominant species tree in which *M. cardinalis* groups with other hummingbird-pollinated taxa (37% of gene trees), rather than being sister to *M. lewisii* (32% of gene trees). We independently corroborate a single evolution of hummingbird pollination syndrome in *Erythranthe* by demonstrating functional redundancy in genetic complementation tests of floral traits in hybrids; together, these analyses overturn a textbook case of pollination-syndrome convergence. Strong asymmetries in allele-sharing (Patterson’s D-statistic and related tests) indicate that gene-tree discordance reflects ancient and recent introgression rather than incomplete lineage sorting. Consistent with abundant introgression blurring the history of divergence, low-recombination and adaptation-associated regions support the new species tree, while high-recombination regions generate phylogenetic evidence for sister status for *M. lewisii* and *M. cardinalis*. Population-level sampling of core taxa also revealed two instances of chloroplast capture, with Sierran *M. lewisii* and Southern Californian *M. parishii* each carrying organelle genomes nested within respective sympatric *M. cardinalis* clades. A recent organellar transfer from *M. cardinalis*, an outcrosser where selfish cytonuclear dynamics are more likely, may account for the unexpected cytoplasmic male sterility effects of selfer *M. parishii* organelles in hybrids with *M. lewisii*. Overall, our phylogenomic results reveal extensive reticulation throughout the evolutionary history of a classic monkeyflower radiation, suggesting that natural selection (re-)assembles and maintains species-diagnostic traits and barriers in the face of gene flow. Our findings further underline the challenges, even in reproductively isolated species, in distinguishing re-use of adaptive alleles from true convergence and emphasize the value of a phylogenomic framework for reconstructing the evolutionary genetics of adaptation and speciation.

**Author Summary:** Adaptive radiations, which involve both divergent evolution of new traits and recurrent trait evolution, provide insight into the processes that generate and maintain organismal diversity. However, rapid radiations also generate particular challenges for inferring the evolutionary history and mechanistic basis of adaptation and speciation, as multiple processes can cause different parts of the genome to have distinct phylogenetic trees. Thus, inferences about the mode and timing of divergence and the causes of parallel trait evolution require a fine-grained understanding of the flow of genomic variation through time. In this study, we used genome-wide sampling of thousands of genes to re-construct the evolutionary histories of a model plant radiation, the monkeyflowers of *Mimulus* section *Erythranthe*. Work over the past half-century has established the parapatric and putatively sister species *M. lewisii* (bee-pollinated) and *M. cardinalis* (hummingbird-pollinated, as are three other species in the section) as textbook examples of both rapid speciation via shifts in pollination syndrome and convergent evolution of floral syndromes. Our phylogenomic analyses re-write both of these stories, placing *M. cardinalis* in a clade with other hummingbird-pollinated taxa and demonstrating that abundant introgression between ancestral lineages as well as in areas of current sympatry contributes to the real (but misleading) affinities between *M. cardinalis* and *M. lewisii*. This work illustrates the pervasive influence of gene flow and introgression during adaptive radiation and speciation, and underlines the necessity of a gene-scale and genome-wide phylogenomics framework for understanding trait divergence, even among well-established species.

## Introduction

Adaptive radiations are engines of biodiversity and thus natural laboratories for understanding its origins [1-5]. During radiations, natural selection can cause both phenotypic divergence as populations move into novel environments and convergence when different populations adapt to similar ecological conditions [6,7]. Divergence provides the opportunity to re-construct the ecological context and genetic basis of adaptive walks, while repeated evolution can reveal the importance of genetic vs. environmental constraints in shaping convergent phenotypes [reviewed in 8]. Furthermore, the processes of adaptation and speciation are tightly intertwined in radiations, and recent radiations help reveal the processes and genes underlying lineage diversification [9-12]. A strong phylogenetic framework is necessary both for understanding the process of speciation and for tracing phenotypic evolution across species (e.g. inferring convergence vs. a single mutational origin for similar phenotypes) [13]. However, the rapid diversification characteristic of adaptive radiations also confounds definition of a single “species tree” [14]. Thus, understanding adaptation and speciation within radiations requires a phylogenomic context that captures the diversity of evolutionary histories across recently diverged genomes [4,15,16].

Two processes confound the reconstruction of a universal genome-wide “species tree”, while also affecting the course of adaptation and speciation [17]. Incomplete lineage sorting (ILS), in which different lineages randomly sample the same alleles polymorphic in their ancestor, can persist after rapid splitting of ancestral populations [18]. In addition, incomplete reproductive isolation between incipient species in areas of sympatry may allow gene flow and introgression that lead to further discordance between the genealogical relationships at any one locus and the deeper species relationships. Both ILS and introgression complicate the inference of species trees, but they have very different impacts on the processes of adaptation. In particular, introgression may cause adaptive alleles, and thus the traits they confer, to be shared among species that are not otherwise closely related [12,19]. Conversely, hybridizing species that are not closely related may appear as sister taxa in phylogenies strongly influenced by introgressed loci (whether those loci are adaptive or not). Such introgression is empirically common, as evidenced by sharp discordance between nuclear and organellar (mitochondrial, chloroplast) phylogenetic trees in many plants [20]and animals [21]. Thus, disentangling the contributions of ILS and introgression to the flow of genetic variation through radiations is important not only to properly characterize the historical process of adaptive evolution, but to reveal its mechanisms. Applying phylogenomic approaches across entire radiations can provide nuanced insight into the constraints, causes, and consequences of adaptive evolution, as well as the processes that structure sequence evolution across complex genomes.

Here, we present phylogenomic re-assessment of the evolutionary history of a classic adaptive radiation in flowering plants, the monkeyflowers of *Mimulus* section *Erythranthe* [22,23]. [Note: Many *Mimulus*, including these taxa, have been re-named as genus *Erythranthe* [24], and the species within this section have been split [25]. However, in the absence of a well-resolved family-level phylogeny, and for consistency with previous work [see 26], we refer to these taxa as *Mimulus* section *Erythranthe* and retain previous species names [23]]. The *Erythranthe* section contains five taxa with flowers adapted for hummingbird pollination (narrow red corolla tubes with little or no landing pad for bees, often abundant nectar; Fig 1). *Mimulus cardinalis* is common in riparian habitats across a broad latitudinal range in western North America (Baja California to Oregon), with disjunct populations occurring in Arizona. The other four hummingbird taxa (*M. eastwoodiae, M. rupestris, M. verbenaceus, M. nelsonii*) are each restricted to much smaller “sky-island” ranges in the southwestern U.S. and Mexico [22,23,25]. The bumblebee-pollinated high-elevation specialist *M. lewisii* is also widespread, with a dark-pink flowered Northern race found in the Rocky and Cascade Mountain ranges [retained as *E. lewisii* in 25] and a pale-pink flowered Sierran race broadly parapatric with *M. cardinalis* in the Sierra Nevada Mountains of California [renamed E. erubescens in 25]. Both the hummingbird- and bee-pollinated taxa are primarily perennial, occurring in soils that remain wet throughout the summer growing season. The eighth taxon, *M. parishii*, is a routinely self-pollinating small-flowered annual occurring in seasonally wet habitats in southern California (e.g. desert washes). Despite their distinct pollination syndromes, all these taxa are at least partially cross-compatible [27,28] and natural hybrids have been reported between *M. cardinalis* and the two taxa with which it co-occurs in California (*M. lewisii* and *M. parishii*) [29]. The combination of diversity and genetic tractability has made the *Erythanthe* radiation a model for understanding the genetic basis of both floral trait divergence and species barriers for over half a century [27].

**Fig 1.**
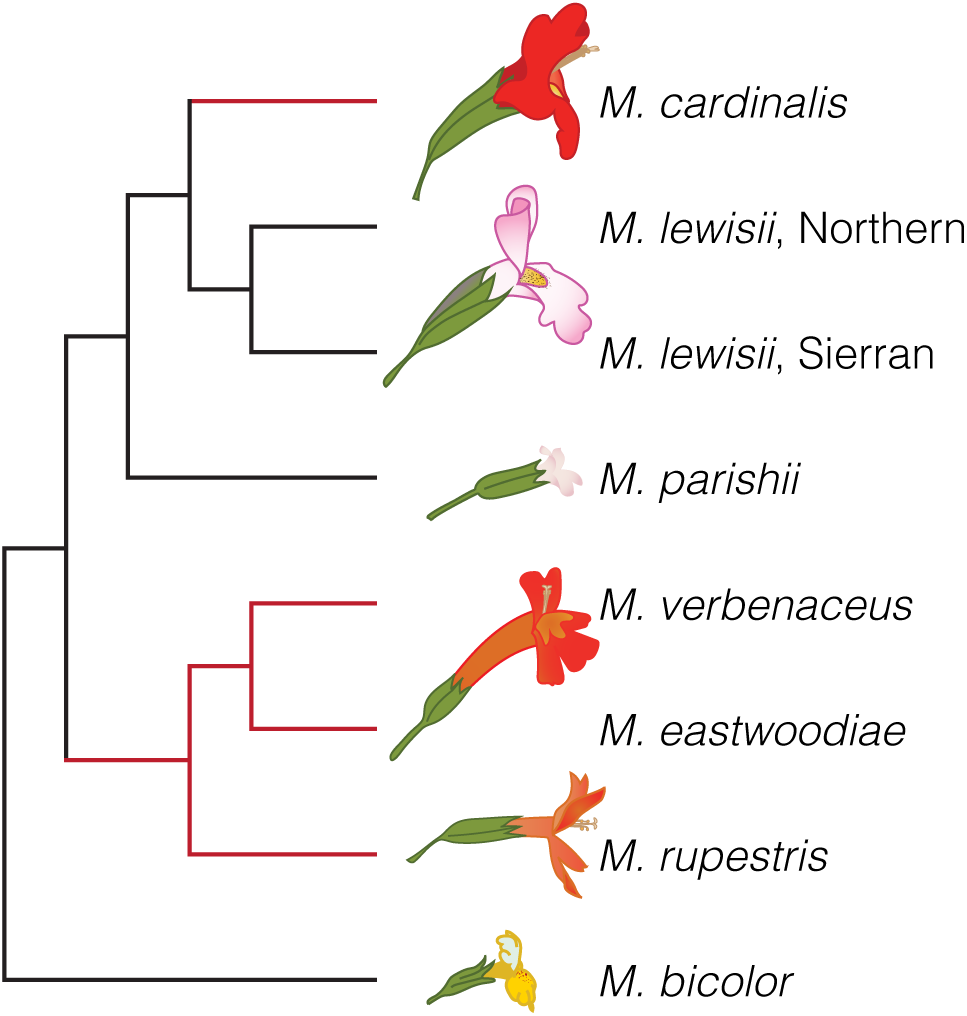
*Mimulus* section *Erythranthe*, with *M. bicolor* as an outgroup, as defined by previous phylogenetic treatments. [22,23,41]. The two putative derivations of hummingbird pollination shown in red.

In ecological genetic work prior to the establishment of molecular phylogenetics, the extensive range overlap and relatively high cross-compatibility Sierran *M. lewisii* and *M. cardinalis* established them as sister taxa locally adapted to distinct elevational and pollinator niches [27,28,30-32]. Groundbreaking QTL mapping studies of species differences and barriers identified the few major loci underlying each aspect of their pollination syndromes, including nectar volume and corolla traits [30,31], and demonstrated that these conferred pollinator specificity and assortative mating between experimental hybrids in sympatry [29,32,33]. It has since become clear that inferring the genetic architecture of adaptation in this pair is complicated by multiple inversions and translocations that suppress free recombination in hybrids [34,35] and also cause underdominant F_1_ sterility [36]. However, the inference that major Mendelian genes define and isolate florally-distinct sister monkeyflowers has been strengthened by the molecular dissection of loci underlying pigmentation variants [37,38], contributing to establishment of this group a model system for floral evolution and development [reviewed in 39]

Sister status for parapatric *M. cardinalis* and *M. lewisii*, and the companion inference of two distinct evolutionary transitions from bee to hummingbird pollination (one in the four sky-island taxa, one more recently in *M. cardinalis*; Fig 1) have remained well-accepted in the post-phylogenetic era. Indeed, after phylogenetic work redefining *Mimulus* [40], re-organizing the North American sections of the genus [41] and re-tracing the evolution of hummingbird pollination in section *Erythranthe* [23], the system became a textbook example of rapid convergent evolution, as well as speciation by large-effect adaptive alleles [e.g. 42]. However, due to low resolution in universal loci used for plant phylogenetics at the time [41], the within-*Erythranthe* tree was primarily based on genome-wide population genetic markers (amplified fragment length polymorphisms, AFLPs) [23]. There are many reasons why either a few slowly-evolving loci or an aggregate of AFLPs might not clearly reflect the true evolutionary history of a given set of species, especially in a recent radiation [15]. Furthermore, while the hummingbird pollination syndrome is one of the most distinct, repeatable, and reproductively-isolating peaks in the adaptive landscape of flowering plants [43-47], inference about the genetic mechanisms of convergence and divergence in pollination syndrome among close relatives requires a well-resolved phylogenetic context. Thus, phylogenomic re-assessment of this group is an essential foundation for the study of micro- and macro-evolutionary processes in this classic system, as well as a window into the complex evolutionary histories possible in even a small radiation.

## Results and Discussion

### Whole-genome species trees suggest a single origin of hummingbird pollination

We used Illumina sequencing of targeted genic regions (gene-capture; see Methods) to survey genome-wide variation within and among species in *Mimulus* section *Erythranthe*. The capture probes targeted genes 1:1 orthologous among *M. lewisii (v 1*.*1; [23]*), M. cardinalis (v 1*.*1*; www.mimubase.org), and the yellow monkeyflower *M. guttatus (*v2 reference; www.Phytozome.jgi.doe.gov)*. We sequenced accessions of *M. lewisii* (n = 19), *M. cardinalis* (n = 34), and *M. parishii* (n =2) from across their geographic ranges, as well as a single accession each of *M. verbenaceus, M. rupestris*, and *M. eastwoodiae* (S1 Table). Across 8,151 sequenced capture regions (7,078,270 bp total) aligned to chromosomes of the v 1.9 *M. cardinalis* reference genome assembly (www.mimulubase.org), we obtained 533,649 single nucleotide variants (SNVs). The bee-pollinated annual *Mimulus bicolor* was used as a close outgroup to section *Erythranthe* [23]. Whole-genome pooled population sequencing of *M. bicolor* revealed an additional 207,238 SNVs between *M. bicolor* and section *Erythranthe* within regions defined by the targeted capture sequencing, totaling 740,887 variant sites. This set of SNVs was divided across 8,151 capture regions with at least one informative site (median: 67 variable sites; IQR: 42-100; max: 316) and fully spans the physical and genetic landscape of *Mimulus* section *Erythranthe* chromosomes, thus providing a well-resolved picture of their evolutionary history.

We inferred phylogenetic relationships among species in Section *Erythranthe* using maximum likelihood inference of the full dataset using IQ-TREE [48] and by assessing variation in gene tree topologies under the multispecies coalescent (MSC) with the software ASTRAL III [49]. Both methods produced identical species relationships (Fig 2, S1 Fig). All species-level branches had 100% bootstrap support (IQ-TREE) and local posterior probabilities of 1 (ASTRAL). ASTRAL quartet scores (i.e. the proportion of underlying gene trees that support a branch in the species tree) ranged from 37.4 to 74.0. Branches closer to our inferred root tended to have lower quartet scores, meaning that a smaller proportion of individual gene trees supported these branches. We interpret the high level of discordance between the species tree and individual gene trees on highly supported branches as the combined effect of ILS and introgression (see below) during the early divergence of ancestral populations.

**Fig 2.**
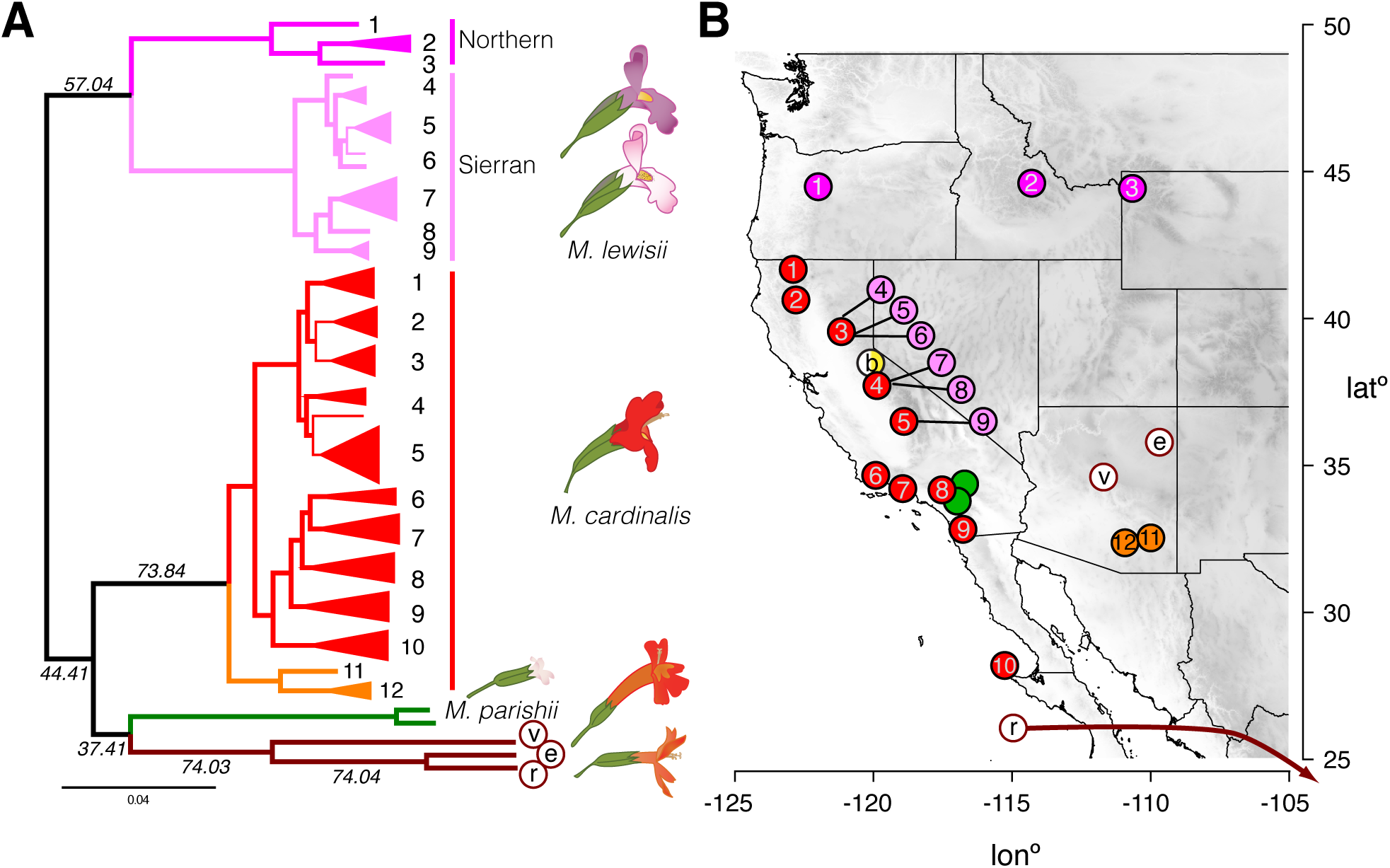
Genome-wide phylogeny of *Mimulus* section *Erythranthe* reveals a single clade containing all hummingbird-pollinated species. (A) The maximum likelihood phylogeny of section *Erythranthe* rooted to *M. bicolor*. The species level topology is identical to that inferred with ASTRAL 3. Branches with bootstrap support >90% are bold. Quartet scores are also given for branches included in the ASTRAL species tree. Clades representing a single collection location are collapsed (see supplemental Fig S8 for the unrooted phylogeny including the branch to *M. bicolor*). Numbers next to *M. lewisii* and *M. cardinalis* tips refer to collection locations in B. (B) Collections of section Erythranthe across the American West. *Sierran M. lewisii* collections are offset due to close overlap with *M. cardinalis* collections in the Sierra Nevada Range. Location of the *M. rupestris* accession from Central Mexico not shown. ‘b’: *M. bicolor*; v: *M. verbenaceus*; ‘e’: *M. eastwoodiae*; ‘r’: *M. rupestris*.

Phylogenetic and phylogeographic patterns within and between *Mimulus lewisii* and *M. cardinalis* are particularly important, given their status as a model system for understanding speciation. Each species formed a monophyletic clade with 100% bootstrap support and phylogeny strongly reflected geography within each. We find a deep split between *M. lewisii* from the Sierra Nevada Range in California (Sierran *M. lewisii; E. erubescens*) and *M. lewisii* from the northern Cascade Range and Rocky Mountains (Northern *M. lewisii; E. lewisii*). This result supports the long-held designation of these two clades as ‘races’ [27] or species [25] (hereafter ‘clades’), based on disjunct ranges, distinct vegetative and floral characters, and partial incompatibility and sterility in some hybrid crosses. *M. cardinalis* was also structured geographically, with accessions from Arizona [named E. *cinnabarina* in 25] forming an outgroup to *M. cardinalis* from the Pacific coast. Within the Pacific clade, *M. cardinalis* from southern California and northern Baja California were monophyletic and sister to a clade containing *M. cardinalis* from the Sierra Nevada. Consistent with the trees, genetic diversity within *M. lewisii* was heavily structured between Northern and Sierran *M. lewisii* (median *d*_*XY*_: 0.0117, IQR: 0.0074-0.0170), and Northern *M. lewisii* was substantially more diverse (median π: 0.0037; IQR: 0.0016-0.0071) than *M. lewisii* in the Sierra Nevada Range (median π: 0.0015; IQR: 0.0006-0.0042). *M. cardinalis* had levels of nucleotide diversity (median π: 0.0036; IQR: 0.0021-0.0060) similar to Northern *M. lewisii* and was more divergent from Sierran *M. lewisii* (median *d*_*XY*_: 0.0151, IQR: 0.0103-0.0203) than the populations of *M. lewisii* were from each other. Observed heterozygosity in *M. cardinalis* decreased with latitude, supporting the hypothesis that the current range of *M. cardinalis* is the result of a recent northward expansion [50]. Additional work will be necessary to determine whether the geographical isolates of both *M. lewisii* and *M. cardinalis* represent fully-fledged species. Regardless, these phenotypically subtle geographic clades make *Erythranthe* an interesting model system for understanding the evolution of postzygotic barriers in allopatry, as well as for the radiation of traits involved in pre-mating isolation in sympatry.

Despite within-species consistency with the previous section *Erythranthe* phylogeny [23], our species tree differs radically in the placement of *M. cardinalis* and *M. parishii*: both are included in a single clade which also contains all other hummingbird-pollinated species (hereafter referred to as Clade H) (Fig 2). The implications for this revision are three-fold. First, the early history of section *Erythranthe* is primarily defined by the split between the ancestor of *M. lewisii* and the common ancestor of all other species in the group. Second, the model pair of *M. lewisii* and *M. cardinalis* do not share recent common ancestry, at least not to the exclusion of any other species in the section. Third, the placement of all red-flowered species in a single clade strongly suggests that the hummingbird pollination syndrome evolved only once in this group and thus is not a case of phenotypic convergence. We therefore address three further questions raised by this inference and its contrast to previous work. Do key hummingbird-associated floral traits in *M. cardinalis* and other red-flowered species share a functional basis? What is the genomic evidence for and against close evolutionary relationships between *M. cardinalis, M. lewisii*, and *M. parishii*? What evolutionary processes are responsible for cross-genome heterogeneity of gene trees in this recent radiation?

### Floral traits in red-flowered species share a functional genetic basis, consistent with a single evolutionary origin

To further investigate whether *M. cardinalis* and the sky-island endemics plausibly share a functional basis for floral traits associated with hummingbird pollination, we conducted a classic genetic complementation test (see Methods). Key hummingbird syndrome traits of both *M. cardinalis* [30,31,34] and the sky-island taxa (e.g. *M. rupestris*) are recessive to *M. lewisii* (as well as *M. parishii;* data not shown), with F_1_ hybrids between bee and hummingbird taxa remarkably *M. lewisii*-like in all floral traits (S2A Fig). Under the historical scenario of convergent evolution from an ancestor resembling bee-pollinated *M. lewisii*, the recessive alleles conferring the hummingbird-associated trait shift (e.g. long styles and anthers, carotenoid pigment) would be independent mutations fixed in each lineage. Thus, unless each series of mutations non-functionalized the same set of target genes, we would expect transgressive segregation in hybrids between the putatively convergent hummingbird taxa. That is, if a causal *a* allele for carotenoid production in *M. cardinalis* (*aaBB*) is not allelic (functionally interchangeable or identical by descent) with the independent *b* allele underlying the phenotype in another taxon (e.g., *M. rupestris* or *M. verbenaceus; AAbb*) the recessive carotenoid phenotype should be masked in F_1_ hybrids (*AaBb*). We see precisely the opposite—the flowers of both F_1_ and F_2_ hybrids between *M. cardinalis* and *M. rupestris* resemble the parents in all respects, with no transgressive *M. lewisii*-like variation (S2B Fig). Divergence between the hummingbird-pollinated species in their floral shape and size leads to segregation beyond parental and F_1_ values in F_2_s, but there is no evidence of hybrids reverting to the dominant *M. lewisii*-like phenotype expected if the genetic basis for the syndrome is not shared. Redundant loss-of-function mutations or epistatic interactions in highly constrained pigmentation pathways could plausibly produce these patterns for corolla color [44]; however, the complementation of the overall floral morphology is best explained by allelism of multiple mutations underlying the shared aspects of the hummingbird pollination syndrome. This independent line of evidence reinforces the phylogenetic inference that the hummingbird pollination syndrome evolved in the common ancestor of *M. cardinalis* and the sky-island endemics, erasing a classic case of convergence and providing a new framework for understanding adaptation and speciation in this model group.

Together, our genomic and experimental results underline the necessity of an explicitly phylogenomic context for understanding trait evolution and speciation in rapid radiations. Hummingbird pollination undoubtedly evolves convergently both within [51] and among [2,43,52] genera, but pollination syndromes may be particularly prone to complex evolutionary histories that mimic phenotypic convergence at low phylogenetic resolution. Like anti-predator mimicry phenotypes in *Heliconius* butterflies [12], specialized pollination syndromes (e.g., hummingbird, moth) evolve to match a pre-existing model [53]. This creates alternative multi-dimensional adaptive peaks separated by valleys of low fitness, although self-pollination may flatten this landscape [54]. Thus, the path from bee to hummingbird pollination appears to be a very narrow and sequential one – that is, a red-flowered mutant without the expected nectar reward or reproductive parts long enough for effective hummingbird pollination may be a poor match for any pollinator [32,55]. Importantly, an intrinsically jagged adaptive landscape may also mean that the joint introgression of multiple traits or their joint retention in the face of homogenizing gene flow (as inferred here) may be common whenever gene exchange occurs during floral diversification. Both processes may mimic true convergence at a coarse phylogenetic scale, but more resemble the repeated re-use of ancient alleles during freshwater adaptation in stickleback populations [56]. As phylogenomic approaches increasingly allow gene-scale investigation of deeper radiations, and more adaptive genes are identified, such sharing of old variation may often be revealed to underlie trait diversification and parallelism, even in otherwise well-resolved species [19,57].

Given the revision of the species tree, it is also worth revisiting the inference that bee-pollination is ancestral [23], especially given the presence of yellow carotenoid pigments in both outgroup taxa such as (bee-pollinated) *M. bicolor* and the hummingbird-pollinated *Erythranthe*. Across flowering plants, transitions from bee to hummingbird pollination appear far more likely than the reverse [47], due either to genetic constraints [51] and/or the ecology of pollination [55]. Bees tend to ignore red flowers and have nowhere to land on narrowly tubular and reflexed “hummingbird” corollas whereas hummingbirds often visit classic bumblebee flowers; for example, hummingbirds made nearly 20% of the visits to Sierran *M. lewisii* in experimental arrays with *M. cardinalis* and hybrids [33]. Even a low frequency of “mistakes”, especially when hummingbird visits are abundant and bees rare, may select for hummingbird-specialization through increased reward, greater attraction, and more precise pollen placement. In this system, where the bee-specialized pale pink flowers and scent production of Sierran *M. lewisii* (*E. erubecens*) appear locally derived [38,58,59], it is plausible that hummingbird visitation to a less-specialized Northern *M. lewisii*-like ancestor precipitating the origin of hummingbird pollination within Clade H. However, ancestral hummingbird pollination remains formally possible and confirming the expected directionality will require reconstruction of the mutational changes contributing to key trait transitions across the entire radiation.

### Extensive introgression creates the evidence for a sister relationship between *M. lewisii* and *M. cardinalis*

Because they are a decades-old model system for understanding the role of reproductive adaptation in plant speciation, general inferences about the nature of those processes hinge on *M. cardinalis* and *M. lewisii* being parapatric sister species. Moreover, the initial inference of a close relationship was plausibly based on similar vegetative morphology, shared geography, and higher genetic compatibility between the Sierran pair than geographically-disjunct populations within each species [27], as well as previous phylogenetic reconstructions [23]. Given that our whole-genome species tree robustly rejects close sister status for *M. lewisii* and *M. cardinalis*, placing *M. cardinalis* within the predominantly hummingbird-pollinated Clade H, it is important to understand the origins of these confounding affinities. Therefore, we examine our genomic dataset for evidence of a close relationship, describe the genomic distribution of regions showing a sister relationship, and infer the processes underlying patterns of gene tree vs. species tree discordance. We used TWISST [60], which quantifies support for different species tree topologies among a set of inferred gene trees, to compare support for trees containing Clade H (all red-flowered species, the ‘species tree’; Fig 3A, orange) to support for trees where *M. lewisii* and *M. cardinalis* form an exclusive clade (the ‘lew-card tree’; Fig 3A, purple). Because we were primarily interested in the relationships between these two focal species, we were agnostic to the placement of *M. parishii* in these analyses. Notably, the lew-card tree was the second-most common topology observed across the genome, next only to our inferred species tree (Fig 3A). Across the entire dataset consisting of 8,151 gene trees, 37% of subtrees identified in TWISST supported the species tree while 32% supported the ‘lew-card’ tree. Substantial incomplete lineage sorting (ILS) at the base of this radiation could produce this pattern, but we hypothesized that introgression between *M. lewisii* and *M. cardinalis* was a more likely source given current parapatry and cross-compatibility. Therefore, to explore introgression as source of gene-tree/species-tree discordance, we tested for (1) asymmetries in patterns of shared, discordant allelic states among species, (2) patterns of absolute genetic divergence indicative of a reticulate evolutionary history, and (3) a correlation between recombination rate and support for the ‘lew-card’ tree.

**Fig 3.**
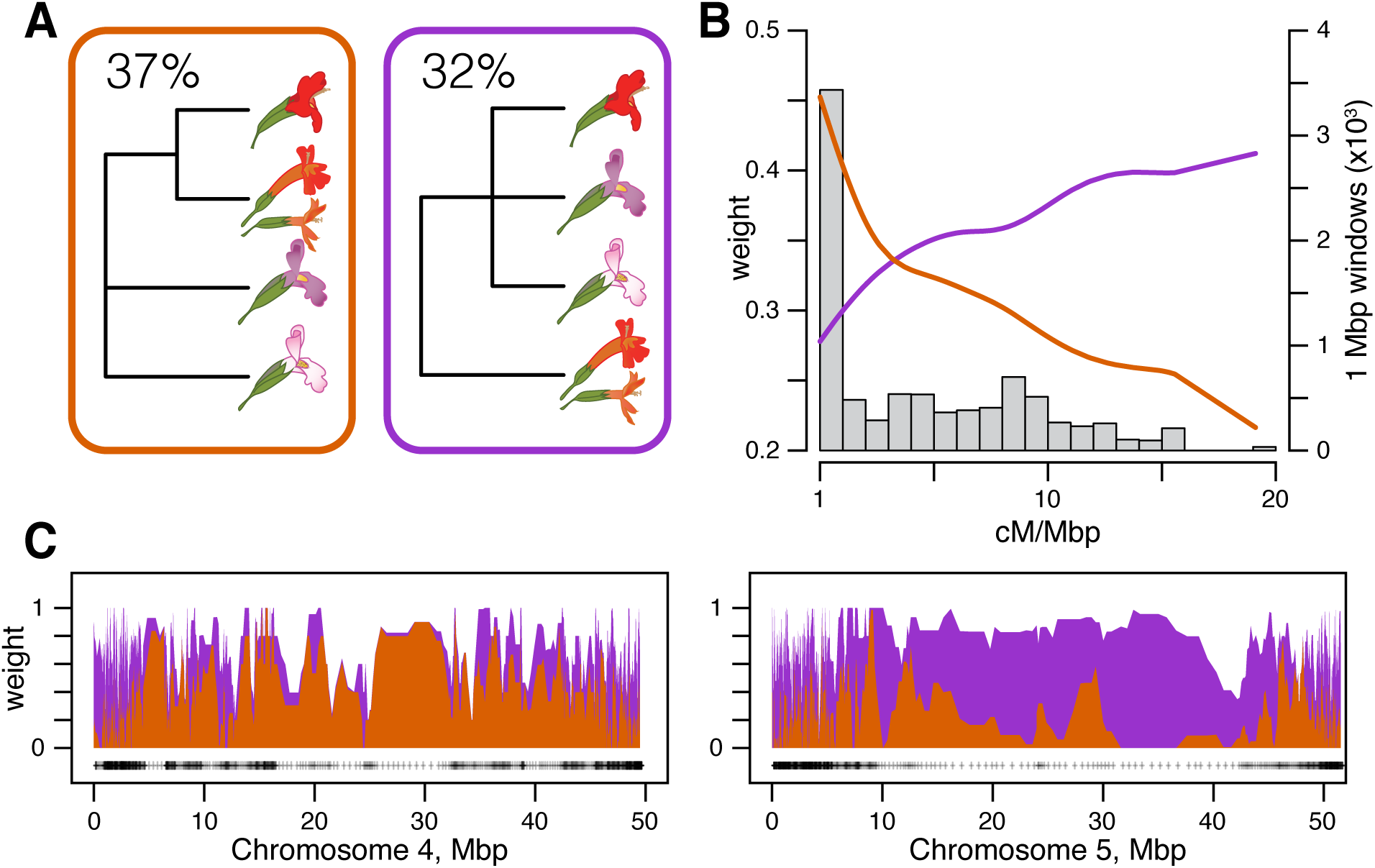
Introgression has generated the evidence for sisterhood of *M. lewisii* and *M. cardinalis*. (A) Genome-wide TWISST weightings for a simplified species tree (orange) and a simplified “lew-card” tree (purple). (B) Support for the species and lew-card trees as a function of recombination rate. Lines show cubic spline fits colored as in A. The gray histogram shows the frequency of genomic windows at a given recombination rate (bin size: 1 cM/Mbp). (C) Topology weights along *M. cardinalis* Chromosomes 4 and 5. Polygons are stacked so that weights across all possible topologies sum to 1. Weights are averaged in windows of 5 genes; black crosses show locations of window midpoints.

We first tested for genome-wide evidence of the presence, timing, and direction of introgression between *M. lewisii* and *M. cardinalis* using Patterson’s D statistic [61] and D_FOIL_ [a five-taxon expansion of Patterson’s D; 62]. Patterson’s D (also known as the ABBA-BABA test) detected significant introgression between *M. cardinalis* and *M. lewisii* (block jackknife: z-score = 3150.844; *p* ∼ 0). The absolute value of D depended on which accessions of *M. cardinalis* and *M. lewisii* were used in the test (range: 0.01 - 0.10), but D was always non-zero (S3 Fig), indicating that introgression was not restricted to a single portion of the current species ranges. Bolstering this inference, the predominant introgression signal detected by D_FOIL_ was between *M. cardinalis* and ancestral *M. lewisii* (i.e., prior to divergence of its Sierran and Northern clades) (S4A Fig). Because the early timing of inferred introgression prevents assessment of its direction with D_FOIL_ alone [62], we used an additional test, D2 [63], which infers the direction of introgression using expectations from the multispecies network coalescent. Directional introgression from *M. cardinalis* into *M. lewisii* would result in reduced nucleotide divergence between *M. lewisii* and the other species of Clade H (e.g. *M. verbenaceus*) at genes following the introgression tree (S4C,D Fig). This is because these alleles sampled from *M. lewisii* are historically *M. cardinalis* alleles and reflect divergence between *M. cardinalis* and the rest of Clade H. In contrast, introgression from *M. lewisii* into *M. cardinalis* would not affect sequence divergence between *M. lewisii* and non-*cardinalis* members of Clade H. We detected no difference in sequence divergence between *M. lewisii* and third taxon *M. verbenaceus* at genes whose history matched the species tree versus the introgression tree (t-test: *t*_3354.3_=1.12, p = 0.26; S4D Fig). Therefore, we infer that introgression during this early period mostly moved genetic material asymmetrically from ancestral *M. lewisii* into *M. cardinalis*.

In addition to producing asymmetric allele-sharing on a phylogeny, the distribution of introgressed DNA should vary predictably across the genome. In particular, the extent to which neutral introgressed variation establishes or fixes in a recipient population should be strongly affected by the local recombination rate [reviewed in 16]. At one extreme, adaptive (or selfish) introgression of a mitochondrial sequence variant could carry both the entire mitochondrial genome and linked chloroplast variants to fixation across species boundaries [64]. However, the more plausible assumption is that the vast majority of genomic segments carry variants that are either neutral or deleterious in a heterospecific background. Because low recombination rates extend the effects of selection against deleterious incoming alleles over larger physical regions, such regions may be broadly protected from introgression. In contrast, variants in high-recombination regions are affected by selection on their individual merits, allowing rates of (neutral or beneficial) introgression to be higher.

To investigate the relationship between recombination rate and introgression in the *Erythranthe* group, we used a dense linkage map of *M. cardinalis* generated from a subset of gene-capture loci [65]. This map supported a chromosome-level scaffolding of *M. cardinalis and M. lewisii* genomes (since reinforced with additional data to form the current V2 genomes; www.mimubase.org) and allows confident genetic-physical comparisons (see Methods). Crossovers in *M. cardinalis* occur almost exclusively on the ends of each chromosome, with very little recombination across large, presumably centromeric and pericentromeric, central regions (S5 Fig). The species tree was the most common topology observed in these low-or non-recombining regions, which also covered ∼68% the physical expanse of the genome (i.e. contigs scaffolded with the genetic map; 235/345 1Mb windows; Fig 3B; S6 Fig). Support for the lew-card tree was strongly and positively correlated with recombination rate (Spearman’s ρ = 0.136, p= 1×10^−10^; Fig 3B), with the introgression topology becoming predominant at recombination rates > 2.5 cM/Mb. Indeed, this pattern is so pervasive that when we inferred the maximum likelihood phylogeny using only SNVs in windows with recombination rates greater than 5 cM/Mb, *M. lewisii* and *M. cardinalis* came out as sister taxa with 100% bootstrap support (S7 Fig). Although elevated introgression only at chromosome ends was the dominant genome-wide pattern, we also observed near-complete replacement of some chromosomes that erased the underlying species tree (Fig 3C,D; S6 Fig). For example, Chromosome 5 consistently supports the ‘lew-card’ tree, including across its low-recombination central region (Fig 3C). In contrast, Chromosome 4 generally showed high support for the species tree (Fig 3C). Chromosome 4 contains multiple ecologically-relevant quantitative trait loci (QTLs) in crosses between *M. lewisii* and *M. cardinalis*, including the ‘yellow upper’ (YUP) locus [30], which switches petal color from pink/purple to red via carotenoid deposition. YUP is embedded in a large region of completely suppressed recombination in *M. lewisii* x *M. cardinalis* mapping crosses (likely an inversion), in tight linkage with a major flower length QTL and a putative hybrid lethality factor [34]. Strong selection against heterospecific alleles and low recombination in hybrids may make this entire chromosome particularly resistant to introgression in areas of ancestral or recent contact between *M. lewisii* and *M. cardinalis*.

Our results corroborate one of most striking results of speciation genomics over the past decade: introgression between closely related species is widespread and can profoundly affect the course of evolution. The extent of introgression ranges from one or a few loci involved in adaptation [12,66] to genome-wide exchange that nearly swamps out past population histories [67-69]. Our phylogenomic results place introgression between *M. lewisii* and *M. cardinalis* near the upper end of this continuum, so it is not surprising that past sampling of loci could infer other histories [23]. Similar patterns have been seen in *Anopheles* mosquitoes [68] and among some cat species [69], where the predominant genome-wide signal derives from hybridization. In those animal cases, strong hybrid F_1_ incompatibilities map to the sex chromosomes, giving them extra weight in inferring the likely species tree. Here, we resolve speciation histories only because these *Mimulus* genomes contain large pericentromeric regions that rarely recombine and are generally resistant to gene exchange. The resulting species-tree inference is bolstered by a strong chromosome-scale match from a key adaptive chromosome (Chromosome 4) underlying multiple pollination-syndrome traits. Within the physically small, but highly recombining and gene-dense ends of chromosomes, admixture predominates. The latter pattern strongly supports our inference that introgression, rather than a recent split, creates signals of sisterhood between *M. lewisii* and *M. cardinalis*.

### Despite strong reproductive barriers between *M. cardinalis* and *M. lewisii*, recent introgression (including chloroplast capture) has occurred in their shared Sierran range

Although broadly parapatric, Sierran *Mimulus lewisii* and *M. cardinalis* are reproductively isolated from one another by a series of strong but incomplete barriers [27,29]. Ecogeographic isolation [29], elevational specialization [70] and distinct pollination syndromes [32] result in near-complete pre-mating isolation. In addition, a pair of intrinsically underdominant chromosomal translocations make F_1_ hybrids >65% pollen-sterile [34,36]. Despite these strong contemporary barriers, we also find substantial evidence of recent introgression (both nuclear and organellar) where *M. lewisii* and *M. cardinalis* co-occur in the Sierra Nevada Range of California. Sierran *M. lewisii* and *M. cardinalis* formed a monophyletic clade in 14.5% of nuclear subtrees analyzed with TWISST; this clade was fully supported at 5.9% of gene trees (479 of 8151). Furthermore, chloroplast haplotypes (genotyped using organellar reads skimmed from the nuclear capture data; see Methods) from Sierran *M. lewisii* and nearby *M. cardinalis* populations form a single clade (100% bootstrap support; Fig 4, S8 Fig). Due to short branch-lengths, we conservatively consider the base of the Sierran *M. lewisii* clade to be a polytomy; however, moderate bootstrap support (62%) for monophyly of the *M. lewisii* haplotypes suggests that a single local *M. cardinalis* cytoplasm may have recently swept through all Sierran *M. lewisii* populations. Importantly, the shared Sierran range where we infer organellar transfer is the source for the accessions of both species used in previous adaptation and speciation genetic studies, phylogenetics [41], and reference genome assemblies.

**Fig 4.**
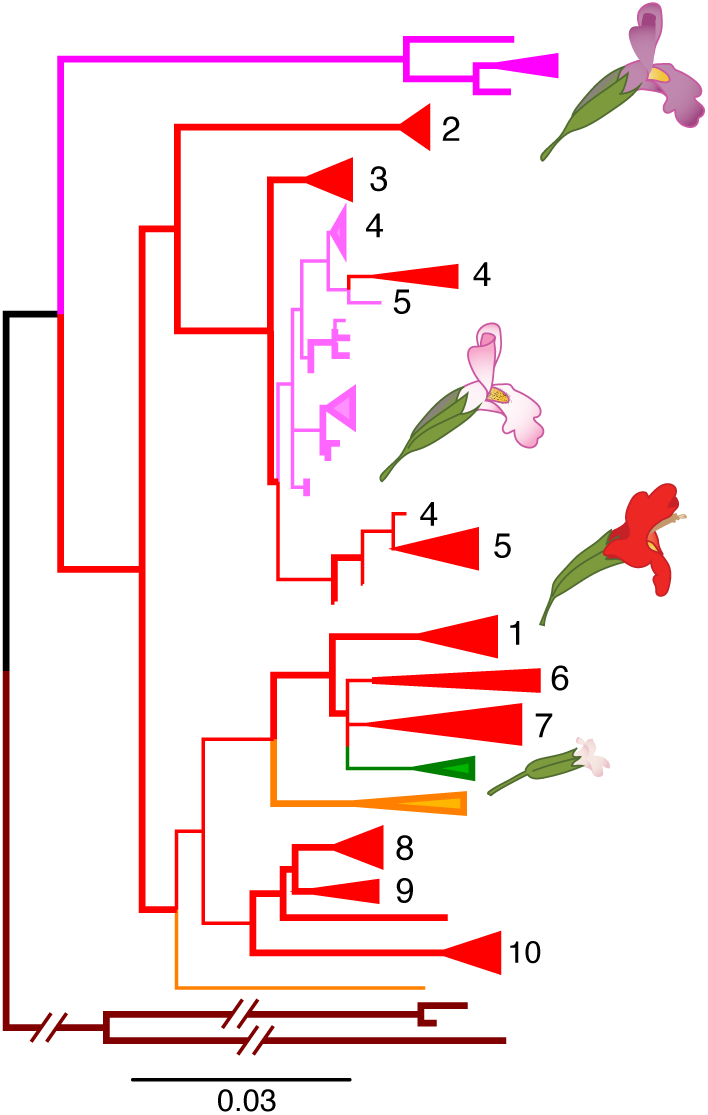
The chloroplast phylogeny demonstrates ancient and recent, geographically local, introgression. The maximum likelihood phylogeny rooted to *M. bicolor* is shown. Long branches to *M. verbenaceus, M. rupestris*, and *M. eastwoodiae* are abbreviated (See S8 Fig for the unrooted, unabbreviated tree). Species and are colored and populations are numbered as in Fig 2. Branches with >90% bootstrap support are in bold.

More work will be necessary to understand whether organellar (and nuclear) introgression in the Sierras represents “surfing” of neutral variation introduced from an expanding *M. cardinalis* range-front [71] or the spread of adaptive or selfish alleles by natural selection. In either case, strong evidence of recent organellar capture [20] reinforces the inference of ancient and recent nuclear introgression in this system, and further suggests that strong ecological and genetic barriers have not been sufficient to isolate the entire genomes of these young taxa upon secondary contact. Although natural hybridization between *M. lewisii* and *M. cardinalis* is rare [29] and costly [29,36], a little gene flow goes a long way [72]. This evidence for recent (as well as ancient) introgression re-iterates the importance of an evolutionary genomic framework for understanding the process of speciation, and also underlines the potential for hybridization (even between highly isolated taxa) as a source of beneficial alleles for contemporary evolution in response to changing environments.

### Organellar capture by selfer *M. parishii* confirms local hybridization with *M. cardinalis*, and may explain cytoplasmic male sterility in its hybrids with *M. lewisii*

In a second case of recent introgression, the chloroplast tree shows that selfing species *M. parishii* has captured the cytoplasmic genomes of the outcrossing *M. cardinalis* (Fig 4). Specifically, *M. parishii* chloroplast haplotypes are nested within *M. cardinalis* variation from their region of range overlap in Southern California. As with the transfer of local *M. cardinalis* organelles into Sierran *M. lewisii*, this geographical signal strongly supports recent introgression over alternative sources of phylogenetic discordance. Despite *M. parishii*’s floral adaptations for self-pollination (tiny pale-pink flowers with little nectar and no separation of male and female organs; Fig 1, Fig S2A), hybrids between the selfer and *M. cardinalis* have been reported where they co-occur along ephemeral waterways. Given the difference in mating system, we might expect that F_1_ hybrids would have selfer seed parents and would backcross primarily to the outcrossing species, causing introgression of nuclear genes from *M. parishii* into *M. cardinalis*, as seen in the yellow monkeyflower pair, *M. nasutus* (selfer) and *M. guttatus* (outcrosser) [73,74]. Instead, the highly selfing species appears to have captured the organellar genome of the outcrossing species. This may have been made more likely by the general dominance of *M. parishii* for floral traits (Fig S2A); in a hybrid swarm, selfing (rather than backcrossing to the outcrossing taxon) may be the primary mode of pollination.

Recent introgression between these highly divergent taxa may also help explain the puzzling cytoplasmic male sterility (CMS; anthers produce no pollen) in hybrids between *M. parishii* and *M. lewisii* [35]. In that study, we found that F_2_ hybrids with the *M. parishii* cytoplasm exhibit CMS if they do not also carry *M. parishii* alleles at multiple nuclear restorer loci, whereas reciprocal hybrids do not exhibit anther sterility. CMS is in flowering plant hybrids is common and thought to result from selfish male-sterilizing mitochondrial haplotypes [75] that spread within species by slightly increasing female fitness, in turn favoring the spread of matched nuclear restorers of male fertility [76]. Selfish CMS-restorer dynamics are theoretically plausible and have been empirically demonstrated in other *Mimulus* species [77], but should not occur in highly selfing taxa where individual female fitness also depends on some pollen production [78]. However, conditions for the spread and establishment of an heterospecific CMS variant, which can co-introgress with its (dominant) restorer allele, may be less restrictive than on a *de novo* CMS mutation. Thus, while *M. parishii* x *M. lewisii* CMS could still reflect independent neutral divergence at the hybrid-interacting loci, *M. parishii*’s possession of an organellar haplotype recently transfered from neighboring *M. cardinalis* revives the possibility of a selfish history for this asymmetric hybrid incompatibility.

## Conclusions

Our understanding of adaption and speciation is contingent on understanding the demographic and genetic histories of diverging populations, which the genomics era is proving to be remarkably reticulate. We present the first population genomic dataset in the classic model system of *Mimulus* section *Erythranthe* to clarify the history of species divergence and reveal rampant introgression during periods of secondary contact. Definitive work on patterns of reproductive isolation [27,29], abiotic [70] and biotic [32]adaptation, convergence in pollination syndromes [23]and speciation genetics [30,36] have been built on the foundation of close sister status for sympatric *M. lewisii* and *M. cardinalis*. However, these model taxa join a growing number of systems in which introgression shapes trait evolution relevant to speciation and obscures deeper histories of divergence. Our analyses suggest that introgressive hybridization – and not recent parapatric speciation – is primarily responsible for the signals of genetic closeness captured in previous phylogenetic analyses (Fig 5). Gene flow between *M. lewisii* and *M. cardinalis*, both in the past and in their current zone of sympatry in the Sierran Nevada Range, causes much of the nuclear genome to support sister species status. Multiple instances of geographically restricted cytoplasmic introgression reinforce the inference of pervasive hybridization in this system and may also explain the paradoxical cytoplasmic male sterility (CMS) of selfer *M. parishii*. Importantly, our revision of the species tree for *Mimulus* section *Erythranthe* demonstrates that long-term resistance to introgression, rather than convergence, may be important in shaping multi-trait pollination syndromes during adaptive radiation in complex landscapes. While shifting the genetic origin of the hummingbird pollination system to an earlier node, our genome-wide evidence for reticulation during the *Erythranthe* radiation only enriches its value for understanding the origins and maintenance of species barriers. The layers of pre- and post-zygotic isolating mechanisms in current contact zones built up over time and space, thus providing the opportunity to excavate their evolution and interactions across the entire radiation.

**Fig 5.**
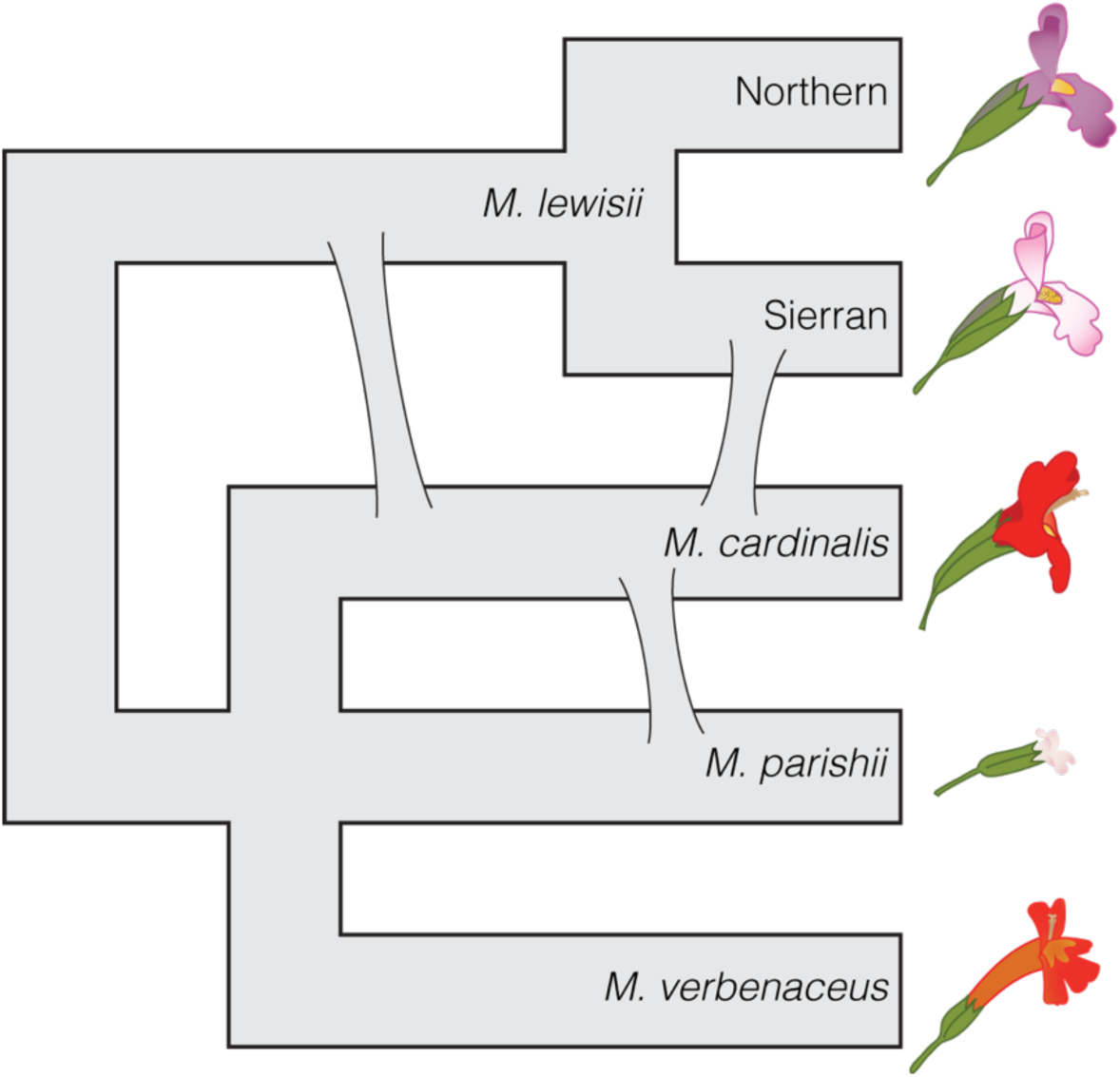
A revised evolutionary history of *Mimulus* section *Erythranthe*. The three major introgression events shown contribute to discordance between previous molecular phylogenies and the revised species tree. The ‘slope’ of each reticulation indicates the inferred direction of introgression. Clade H is shown as a tritomy due to long external branches and short internal branches; however, it is plausible that *M. parishii* and a hummingbird-pollinated ancestor of *M. cardinalis* and *M. verbenaceus* were both separately derived from an (large, potentially structured) ancestral population that phenotypically resembled Northern *M. lewisii*.

## Materials and Methods

### Collections and plant material

We obtained wild-collected seeds from throughout the geographic range of *Mimulus* section *Erythranthe* with particular focus on *M. lewisii* and *M. cardinalis* populations (Fig 2, Suppl. Table S1). Plants were grown from seed in a greenhouse at the University of Montana and DNA extracted from leaf or flower bud tissue using a customized CTAB-chloroform extraction protocol (dx.doi.org/10.17504/protocols.io.bgv6jw9e). We used *M. bicolor* as an outgroup species to the core *Erythranthe* taxa. Whole *M. bicolor* plants (n = 160) were wild-collected from a large color-polymorphic population in center of its range in the Sierra Nevada Range [79] and dried in coin envelopes, and then DNA was extracted from tissue individually prior to equal-volume pooling.

### Linkage mapping and recombination rates

We used the *M. cardinalis* linkage map reported in [65] and CE10 v1.92 genome contigs (www.mimubase.org) to estimate genetic and physical distances along the *M. cardinalis* genome. Briefly, a Sierran (CE10 inbred line) x Southern (WFM) *M. cardinalis* F_2_ mapping population (N = 93) was genotyped using the same targeted capture approach as this study. 8100 snps (representing 2152 cross-informative capture targets) were ordered with Lep-MAP3 [80], resolving the expected 8 linkage groups (2N = 16) spanning 573 centiMorgans (cM) [65]. The linkage map was used to scaffold v1.92 contigs with Chromonomer version 1.08 [81]. We were able to scaffold a total of 341.8 Mb of genome sequence, which is 83.6**%** of the current v2 chromosomal assembly (based on both optical mapping and linkage relationships; www.mimubase.org). The genome scaffolding used here for genome scans is largely similar in order to the v2 assembly, but its contig positions and orientation are based solely on intraspecific recombination. Recombination rates were estimated in non-overlapping genomic bins of 1 Mbp. Rates were calculated as the genetic distance (in cM) between the two most distal markers in the bin divided by their physical distance (in Mbp). We removed three bins with extreme recombination rate estimates (>100 cM/Mbp) from further analysis. These estimates were due to many crossovers between putatively physically proximal markers (<5,000 bp) with no other markers present in the bin, and likely represent mislocalization of a marker on the physical sequence (e.g., due to paralogy).

### Targeted capture sequencing and genotyping

Targeted sequence capture was used to high-coverage, high-quality genotyping within and among species in *Mimulus* section *Erythranthe*. Capture baits were designed to tile 9,126 genes that are 1:1 orthologous between *M. cardinalis, M. lewisii*, and *M. guttatus*. Details of bait design and library preparation can be found in [65]. All libraries were sequenced on a single lane of Illumina HiSeq 2500 (PE 125). Raw Illumina reads were quality filtered and trimmed for sequencing adaptors using Trimmomatic [82] and aligned to the v1.9g draft *M. cardinalis* genome (http://mimubase.org/FTP/Genomes/) using bwa-mem v0.7.15 [83]. Alignments were filtered for minimum quality scores of 29 using samtools v1.3 [84]. We then removed potential PCR duplicates and realigned around indels using Picard Tools (http://broadinstitute.github.io/picard) and GATK (v3.3-0-g37228af) [85] following GATK best practices.

### Pooled population sequencing of *M. bicolor*

*Mimulus bicolor* DNA (N = 160 wild plants from a large population) was pooled into a single sample for this study. Illumina library preparation and sequencing on an Illumina HiSeq 4000 were performed by Novogene Corporation (Stockton, CA, USA) following manufacturer protocols. Genotypes were called as above with the exception of two alterations intended to convert pooled genotypes into a single *M. bicolor* reference alignment. First, during GVCF creation, we instructed the GATK tool HaplotypeCaller to attempt to remove ‘contaminant’ reads at frequencies of up to 10**%** in order to remove low-frequency polymorphisms present in the pool. After VCF creation, we converted remaining heterozygous sites to homozygotes by randomly selecting one of the two alternate alleles. Multi-allelic sites were all ignored in the final analyses. Observed sequence divergence between *M. bicolor* and *M. cardinalis* (median d_xy_: 0.0277) was similar to levels of synonymous site diversity observed within a single population of the genus’s flagship species, *M. guttatus* [86] aligned and genotyped using the similar parameters. Additionally, observed *M. bicolor—M*.*cardinalis* sequence divergence was nearly identical to *M. bicolor—M. lewisii* divergence (median d_xy_: 0.0282). These results indicate that reference bias is of relatively low concern in this largely genic dataset, despite its phylogenetic scope.

### Gene tree and species tree inference

To generate a set of genomic regions representing individual protein-coding genes, we aligned capture bait sequences to the contig-level *M. cardinalis* v 1.9g genome assembly (http://mimubase.org/FTP/Genomes/) using BLAST v2.2.31 [87] to determine the beginning and end coordinates of each aligned bait. We then used bedtools-merge v2.26.0 [88] to merge bait alignments tiling the same gene into a single region, resulting in 8151 genomic regions. Because each capture region was designed to target a protein-coding gene, we refer to these targeted genomic regions as “genes”.

Gene tree inference and partitioned maximum likelihood (ML) phylogenetic analysis were performed on individual alignments representing each gene. We created individual alignments by extracting genotypes within the boundaries of each gene from the phased VCF using tabix [89]. Alignments thus consisted of variable sites only, and a single haplotype for each sample was included. We inferred ML phylogenies for each gene individually and the entire genome using IQ-TREE v1.7-beta14 (cite) [48] under the GTR+ASC+G4 substitution model to correct for the absence of invariant sites. This dataset included 8,151 genes in which we observed parsimony-informative sites. For the whole-genome phylogeny we also generated branch support by performing 1000 ultrafast bootstrap replicates [90]. To further ensure that the resulting phylogeny was robust to model assumptions and tree search strategies, we inferred ML trees using PhyML v20120412 [91] and RAXML v8.2.12 [92] on a concatenated super-matrix consisting of 600,267 variable sites under the GTR+gamma substitution model with four rate categories.

In addition to whole-genome concatenation, we used ASTRAL-III v5.6.3 [49] to generate a species tree under the multispecies coalescent. ASTRAL uses variation in gene tree topologies to infer a species tree under the assumption that topological discordance among gene trees is due to incomplete lineage sorting during population divergence. We ran ASTRAL on the full dataset of 8,151 gene trees inferred from IQ-TREE, using quartet scores and local posterior probabilities as branch supports. Quartet scores measure how often a given quartet (unrooted, four-taxon tree) observed in the species tree is present in the underlying gene trees. Under the assumption of no gene flow post-speciation, quartet scores are also indicative of the degree of incomplete lineage sorting along the inferred branch [93].

### Tree topology weighting with TWISST

We quantified this variation in species relationships throughout the genome using TWISST [60]. Given a gene tree and a set of species designations for all tips in the tree, TWISST quantifies support for all possible (rooted) species trees through iterative sampling of subtrees where each species is represented by a single tip. We ran TWISST on each gene tree grouping all accessions by species except *M. verbenaceus, M. rupestris*, and *M. eastwoodiae*, which we grouped into a single ‘species.’ We did this for three reasons: (1) these species formed a single, highly supported clade in our ML and ASTRAL trees, (2) we were primarily interested in the relationships between *M. lewisii* and *M. cardinalis*, and (3) collapsing these species limited our analysis to five taxa (105 unique rooted trees) and made analysis of the entire dataset feasible (vs. seven taxa: 10,395 unique rooted trees). To quantify support among generalized species relationships (e.g. Fig 3A), topology weightings for each unique tree topology were summed across all topologies that included a clade of interest. For instance, we calculated support for the ‘species tree’ as the sum of weightings across all topologies that place *M. cardinalis* in a clade with the other red-flowered species. We also visualized support for different species relationships across the *M. cardinalis* genome by updating genome coordinates of capture regions to match the chromosome-level v2 reference assembly (www.mimubase.org). To aid in visualization, we averaged topology weights in overlapping five-gene windows.

### Genome-wide tests for introgression

We used Patterson’s D [61] and related statistics to identify aggregate genomic signatures of introgression, assuming our inferred species tree accurately reflects historical relationships within section *Erythranthe*. All tests were implemented in Python v3.5.5. Patterson’s D statistics tested for introgression on the four-taxon tree of (*M. bicolor*, (*M. lewisii*, (*M. cardinalis, M. verbenaceus*))). Calculating D using *M. parishii* instead of *M. verbenaceus* produced qualitatively similar results. We used all pairwise combinations of individual accessions of *M. lewisii* and *M. cardinalis*, allowing for heterozygosity but not missing data. While D can be calculated from allele frequencies, our accessions represent multiple populations that may have experienced variable histories of introgression; pairwise calculation gave us the potential to detect geographically-limited introgression. To test for genome-wide statistical significance, we implemented the genomic window jackknife procedure suggested in [94].

D_FOIL_ statistics [62] were used to identify the timing and, potentially, the direction of introgression on the five-taxon tree (*M. bicolor*, ((*M. verbenaceus, M. cardinalis*), (Sierran *lewisii*, Northern *lewisii*))). As with Patterson’s D, we implemented D_FOIL_ in Python using individual accessions and allowing for heterozygosity but not missing data. Because the D_FOIL_ patterns we observed prevented us from inferring the direction of introgression, we calculated Hahn and Hibbins’ D2 [63]. D2 uses expectations from the network coalescent to infer the direction of introgression on a three-taxon tree. We defined the species tree as ((*M. verbenaceus, M. cardinalis*), *M. lewisii*) and the introgression tree as (*M. verbenaceus*, (*M. cardinalis, M. lewisii*)). Introgression from *M. cardinalis* into *M. lewisii* will also result in *M. lewisii* and *M. verbenaceus* sharing more recent common ancestry than at gene trees concordant with the species tree, while introgression from *M. lewisii* into *M. cardinalis* will not. We tested for this difference ([dxy_lew-verb_ | species tree] - [dxy_lew-verb_ | introgression tree]) using a t-test on genes with full TWISST weighting for either the simplified species tree or the simplified introgression tree (see Fig 3).

### Nucleotide diversity and divergence

Population genetic statistics were all calculated with the Python module scikit-allel v1.2.1 https://scikit-allel.readthedocs.io/en/stable/index.html. As input, VCF files were created that included invariant sites using the flag “--includeNonVariantSites” in the GATK tool GenotypeGVCFs. We calculated statistics on our pre-defined capture regions (‘genes’). Nucleotide diversity (π) at each gene was calculated at the species and regional levels (e.g. *M. lewisii* and *Sierran lewisii*) and nucleotide divergence (d_xy_) was calculated among regions and species. In the absence of a complete reference annotation for *M. cardinalis*, we did not differentiate among codon positions or between coding and noncoding diversity.

### Floral trait complementation test

As a rough test for allelism of genetic variation contributing to the hummingbird pollination floral syndrome of *M. cardinalis* and the other red-flowered taxa (specifically *M. verbenaceus* and *M. rupestris*) within the frame of the historical phylogeny, we used a classic complementation approach. First, we generated F_1_ hybrids by crossing *M. rupestris* and *M. verbenaceus* lines (Table S1) to the putative ancestral bee-pollinated phenotype represented by *M. lewisii* (Sierran LF10 line) to verify that these taxa shared recessive inheritance of the hummingbird syndrome phenotype with *M. cardinalis (*. Second, we generated F_1_ hybrids between the CE10 *M. cardinalis* line and *M. rupestris* and *M. verbenaceus*, and then made F_2_s by selfing a single F_1_ of each pair. We grew parents (N= 8-10), F_1_s (N = 10) and F_2_s (N = 100-200) in the greenhouse at the University of Montana. For both sets of hybrids, it was evident that the overall morphology and color of hybrid flowers exhibited non-complementation (Fig S1B). However, severe hybrid breakdown (e.g., deformed corollas, sterile anthers) was also common in both sets of F_2_s. Due to the latter (and the complete absence of obviously *M. lewisii*-like variants), we do not report F_2_ quantitative traits.

## Acknowledgments

We are grateful to Katie Zarn and Jacob Heiling for assistance with plant care, and to Kayli Anderson and Tamara Max and Deng-hui (David) Xing of the University of Montana Genomics Core Facility for assistance with the laboratory work. Mariah McIntosh created the floral illustrations. Amy Angert, Robert Vickrey, Jay Sobel, Thomas Mitchell-Olds, Margaret Hendrick, Seema Sheth, and Brooke Kern generously provided tissue or seeds. We also thank Paul Beardsley and attendees of the Mimulus Meeting 2019 for lively discussions. Funding was provided by NSF DEB-1407333 to AS and LF, and NSF DEB-1457763 and OIA-1736249 to LF. TCN was supported by an UNVEIL postdoctoral fellowship through OIA-1736249.

## Author Contributions

TCN, AMS, and LF conceived of and designed the project. AMS collected wild accessions, prepared tissue, and performed DNA extraction. AMS. FRF, and LF generated and anlayzed M. cardinalis x M. rupestris and M. cardinalis x M. verbenaceus F_2_ populations. AMS and DDV designed the targeted capture probes and oversaw sequence library preparation, in consultation with LF and FRF. YY provided the M. cardinalis draft genome sequence. TCN conducted bioinformatic and phylogenomic analyses, with advice from DDV. TCN and LF wrote the manuscript, with input from all authors.

## Data availability

Code required to generate figures and statistics in the text can be found at github.com/thomnelson/MimulusPhylogenomics. Sequence data will be made publicly available on the SRA.

## References

1. Gavrilets S, Losos JB. Adaptive radiation: contrasting theory with data. Science. 2009;323: 732–737. doi: 10.1126/science.1157966

2. Stebbins GL. Adaptive radiation of reproductive characteristics in angiosperms I: Pollination mechanisms. Annu Rev Ecol Syst. 1970;1: 307–326. doi:https://doi.org/10.1146/annurev.es.01.110170.001515

3. Berner D, Salzburger W. The genomics of organismal diversification illuminated by adaptive radiations. Trends Genet. 2015;31: 491–499. doi: 10.1016/j.tig.2015.07.002

4. Marques DA, Meier JI, Seehausen O. A combinatorial view on speciation and adaptive radiation. Trends Ecol Evol. 2019;34: 531–544. doi: 10.1016/j.tree.2019.02.008

5. Schluter D. The Ecology of Adaptive Radiation. Oxford: Oxford University Press; 2000.

6. Schluter D, Nagel LM. Parallel Speciation by Natural Selection. Am Nat. 1995;146: 292–301. doi: 10.2307/2463062

7. Mahler DL, Ingram T, Revell LJ, Losos JB. Exceptional convergence on the macroevolutionary landscape in island lizard radiations. Science. 2013;341: 292–295. doi: 10.1126/science.1232392

8. Elmer KR, Meyer A. Adaptation in the age of ecological genomics: insights from parallelism and convergence. Trends Ecol Evol. 2011;26: 298–306. doi: 10.1016/j.tree.2011.02.008

9. Seehausen O, Butlin RK, Keller I, Wagner CE, Boughman JW, Hohenlohe PA, et al. Genomics and the origin of species. Nature Rev Genet. 2014;15: 176–192. doi: 10.1038/nrg3644

10. Filiault DL, Ballerini ES, Mandáková T, Aköz G, Derieg NJ, Schmutz J, et al. The *Aquilegia* genome provides insight into adaptive radiation and reveals an extraordinarily polymorphic chromosome with a unique history. Elife. 2018;7: e36426. doi: 10.7554/eLife.36426

11. Meier JI, Marques DA, Wagner CE, Excoffier L, Seehausen O. Genomics of Parallel Ecological Speciation in Lake Victoria Cichlids. Mol Biol Evol. 2018;35: 1489–1506. doi: 10.1093/molbev/msy051

12. Heliconius Genome Consortium. Butterfly genome reveals promiscuous exchange of mimicry adaptations among species. Nature. 2012;487: 94–98. doi: 10.1038/nature11041

13. Glor RE. Phylogenetic insights on adaptive radiation. Annu Rev Ecol Evol Syst. 2010;41: 251–270. doi: 10.1146/annurev.ecolsys.39.110707.173447

14. Nichols R. Gene trees and species trees are not the same. Trends Ecol Evol. 2001;16: 358–364. doi: 10.1016/s0169-5347(01)02203-0

15. Degnan JH, Rosenberg NA. Gene tree discordance, phylogenetic inference and the multispecies coalescent. Trends Ecol Evol. 2009;24: 332–340. doi: 10.1016/j.tree.2009.01.009

16. Payseur BA, Rieseberg LH. A genomic perspective on hybridization and speciation. Mol Ecol. 2016;25: 2337–2360. doi: 10.1111/mec.13557

17. Edwards SV. Is a new and general theory of molecular systematics emerging? Evolution. 2009;63: 1–19. doi: 10.1111/j.1558-5646.2008.00549.x

18. Rosenberg NA, Nordborg M. Genealogical trees, coalescent theory and the analysis of genetic polymorphisms. Nature Rev Genet. 2002;3: 380–390. doi: 10.1038/nrg795

19. Jones MR, Mills LS, Alves PC, Callahan CM, Alves JM, Lafferty DJR, et al. Adaptive introgression underlies polymorphic seasonal camouflage in snowshoe hares. Science. 2018;360: 1355–1358. doi: 10.1126/science.aar5273

20. Rieseberg LH, Soltis DE. Phylogenetic consequences of cytoplasmic gene flow in plants. Evol Trends Plants. 1991;5: 65–84.

21. Toews DPL, Brelsford A. The biogeography of mitochondrial and nuclear discordance in animals. Mol Ecol. 2012;21: 3907–3930. doi: 10.1111/j.1365-294X.2012.05664.x

22. Vickery RK Jr, Wullstein BM. Comparison of six approaches to the classification of *Mimulus* sect. *Erythranthe* (Scrophulariaceae). Syst Bot. 1987;12: 339–364. doi: 10.2307/2419258

23. Beardsley P, Yen A, Olmstead RG. AFLP phylogeny of *Mimulus* section *Erythranthe* and the evolution of hummingbird pollination. Evolution. 2003;57: 1397–1410. doi: 10.1111/j.0014-3820.2003.tb00347.x

24. Barker W, Nesom G, Beardsley P, Fraga NS. A taxonomic conspectus of Phrymaceae: A narrowed circumscription for *Mimulus,* new and resurrected genera, and new names and combinations. Phytoneuron. 2012;39: 1–60.

25. Nesom GL. Taxonomy of *Erythranthe* sect. *Erythranthe* (Phrymaceae). Phytoneuron. 2014;31: 1–41.

26. Lowry DB, Sobel JM, Angert AL, Ashman T-L, Baker RL, Blackman BK, et al. The case for the continued use of the genus name *Mimulus* for all monkeyflowers. Taxon. 2019;68: 617–623. doi: 10.1002/tax.12122

27. Hiesey W, Nobs M, Bjorkman O. Experimental studies on the nature of species: 5. Biosystematics, genetics and physiological ecology of the Erythranthe section of Mimulus. Washington, D.C.: Carnegie Institution; 1971.

28. Vickery RK Jr. Case studies in the evolution of species complexes in *Mimulus*. In: Hecht MK, Steere WC, Wallace B, editors. Evolutionary Biology. 3rd ed. Boston, MA. 1978. pp. 405–507. doi: 10.1007/978-1-4615-6956-5_7

29. Ramsey J, Bradshaw HD Jr., Schemske DW. Components of reproductive isolation between the monkeyflowers *Mimulus lewisii* and *M. cardinalis* (Phrymaceae). Evolution. 2003;57: 1520–1534. doi: 10.1111/j.0014-3820.2003.tb00360.x

30. Bradshaw HD Jr., Wilbert SM, Otto KG, Schemske DW. Genetic mapping of floral traits associated with reproductive isolation in monkeyflowers (*Mimulus*). Nature. 1995;376: 762–765. doi: 10.1038/376762a0

31. Bradshaw HD Jr., Otto KG, Frewen BE, Makowsky R, Schemske DW. Quantitative trait loci affecting differences in floral morphology between two species of monkeyflower (*Mimulus*). Genetics. 1998;149: 367–382.

32. Bradshaw HD Jr., Schemske DW. Allele substitution at a flower colour locus produces a pollinator shift in monkeyflowers. Nature. 2003;426: 176–178. doi: 10.1038/nature02106

33. Schemske DW, Bradshaw HD Jr. Pollinator preference and the evolution of floral traits in monkeyflowers (Mimulus). Proc Nat Acad Sci USA. National Academy of Sciences; 1999;96: 11910–11915. doi: 10.1073/pnas.96.21.11910

34. Fishman L, Stathos A, Beardsley P, Williams CF, Hill JP. Chromosomal rearrangements and the genetics of reproductive barriers in *Mimulus* (monkeyflowers). Evolution. 2013;67: 2547–2560. doi: 10.1111/evo.12154

35. Fishman L, Beardsley P, Stathos A, Williams CF, Hill JP. The genetic architecture of traits associated with the evolution of self-pollination in *Mimulus*. New Phytol. 2015;205: 907–917. doi: 10.1111/nph.13091

36. Stathos A, Fishman L. Chromosomal rearrangements directly cause underdominant F1 pollen sterility in *Mimulus lewisii*–*Mimulus cardinalis* hybrids. Evolution. 2014;68: 3109–3119. doi: 10.1111/evo.12503

37. Yuan Y-W, Rebocho AB, Sagawa JM, Stanley LE, Bradshaw HD Jr. Competition between anthocyanin and flavonol biosynthesis produces spatial pattern variation of floral pigments between *Mimulus* species. Proc Nat Acad Sci USA. 2016;113: 2448–2453. doi: 10.1073/pnas.1515294113

38. Yuan Y-W, Sagawa JM, Young RC, Christensen BJ, Bradshaw HD Jr. Genetic dissection of a major anthocyanin QTL contributing to pollinator-mediated reproductive isolation between sister species of *Mimulus*. Genetics. 2013;194: 255–263. doi: 10.1534/genetics.112.146852

39. Yuan Y-W. Monkeyflowers (Mimulus): new model for plant developmental genetics and evo-devo. New Phytol. 2019;222: 694–700. doi: 10.1111/nph.15560

40. Beardsley P, Olmstead RG. Redefining Phrymaceae: the placement of *Mimulus*, tribe Mimuleae, and *Phryma*. Am J Bot. 2002;89: 1093–1102. doi: 10.3732/ajb.89.7.1093

41. Beardsley P, Schoenig SE, Whittall JB, Olmstead RG. Patterns of evolution in western North American *Mimulus* (Phrymaceae). Am J Bot. 2004;91: 474–489. doi: 10.3732/ajb.91.3.474

42. Freeman S, Herron JC. Evolutionary Analysis. 4 ed. Upper Saddle River, NJ: Pearson/Prentice Hall; 2004.

43. Marais des DL, Rausher MD. Parallel evolution at multiple levels in the origin of hummingbird pollinated flowers in *Ipomoea*. Evolution. John Wiley & Sons, Ltd; 2010;64: 2044–2054. doi: 10.1111/j.1558-5646.2010.00972.x

44. Wessinger CA, Rausher MD. Lessons from flower colour evolution on targets of selection. J Exp Bot. 2012;63: 5741–5749. doi: 10.1093/jxb/ers267

45. Chase MA, Stankowski S, Streisfeld MA. Genomewide variation provides insight into evolutionary relationships in a monkeyflower species complex (*Mimulus* sect. *Diplacus*). Am J Bot. 2017;104: 1510–1521. doi: 10.3732/ajb.1700234

46. Ortiz-Barrientos D. The color genes of speciation in plants. Genetics. 2013;194: 39–42. doi: 10.1534/genetics.113.150466

47. Thomson JD, Wilson P. Explaining evolutionary shifts between bee and hummingbird pollination: convergence, divergence, and directionality. Int J Plant Sci. 2008;169: 23–38. doi: 10.1086/523361

48. Nguyen L-T, Schmidt HA, Haeseler von A, Minh BQ. IQ-TREE: a fast and effective stochastic algorithm for estimating maximum-likelihood phylogenies. Mol Biol Evol. 2015;32: 268–274. doi: 10.1093/molbev/msu300

49. Zhang C, Rabiee M, Sayyari E, Mirarab S. ASTRAL-III: polynomial time species tree reconstruction from partially resolved gene trees. BMC Bioinformatics. 2018;19: 153–30. doi: 10.1186/s12859-018-2129-y

50. Sheth SN, Angert AL. Demographic compensation does not rescue populations at a trailing range edge. Proc Nat Acad Sci USA. 2018;115: 2413–2418. doi: 10.1073/pnas.1715899115

51. Wessinger CA, Rausher MD, Hileman LC. Adaptation to hummingbird pollination is associated with reduced diversification in *Penstemon*. Evol Lett. 2019;3: 521–533. doi: 10.1002/evl3.130

52. Fenster CB, Armbruster WS, Wilson P, Dudash MR, Thomson JD. Pollination syndromes and floral specialization. Annu Rev Ecol Evol Syst. 2004;35: 375–403. doi: 10.1146/annurev.ecolsys.34.011802.132347

53. Bleiweiss R. Mimicry on the QT(L): genetics of speciation in *Mimulus*. Evolution. Society for the Study of Evolution; 2001;55: 1706–1709. doi: 110.1111/j.0014-3820.2001.tb00690.x

54. Wessinger CA, Kelly JK. Selfing can facilitate transitions between pollination syndromes. Am Nat. 2018;191: 582–594. doi: 10.1086/696856

55. Gegear RJ, Burns R, Swoboda-Bhattarai KA. “Hummingbird” floral traits interact synergistically to discourage visitation by bumble bee foragers. Ecology. 2017;98: 489–499. doi: 10.1002/ecy.1661

56. Nelson TC, Cresko WA. Ancient genomic variation underlies repeated ecological adaptation in young stickleback populations. Evol Lett. 2018;2: 9–21. doi: 10.1002/evl3.37

57. Edelman NB, Frandsen PB, Miyagi M, Clavijo B, Davey J, Dikow RB, et al. Genomic architecture and introgression shape a butterfly radiation. Science. 2019;366: 594–599. doi: 10.1126/science.aaw2090

58. Byers KJRP, Vela JP, Peng F, Riffell JA, Bradshaw HD Jr. Floral volatile alleles can contribute to pollinator-mediated reproductive isolation in monkeyflowers (*Mimulus*). Plant J. 2014;80: 1031–1042. doi: 10.1111/tpj.12702

59. Peng F, Byers KJRP, Bradshaw HD Jr. Less is more: Independent loss-of-function OCIMENE SYNTHASE alleles parallel pollination syndrome diversification in monkeyflowers (*Mimulus*). Am J Bot. Wiley-Blackwell; 2017;104: 1055–1059. doi: 10.3732/ajb.1700104

60. Martin SH, Van Belleghem SM. Exploring evolutionary relationships across the genome using topology weighting. Genetics. Genetics; 2017;206: 429–438. doi: 10.1534/genetics.116.194720

61. Durand EY, Patterson N, Reich D, Slatkin M. Testing for ancient admixture between closely related populations. Mol Biol Evol. 2011;28: 2239–2252. doi: 10.1093/molbev/msr048

62. Pease JB, Hahn MW. Detection and polarization of introgression in a five-taxon phylogeny. Syst Biol. 2015;64: 651–662. doi: 10.1093/sysbio/syv023

63. Hahn MW, Hibbins MS. A three-sample test for introgression. Mol Biol Evol. 2019;36: 2878–2882. doi: 10.1093/molbev/msz178

64. Sloan DB, Havird JC, Sharbrough J. The on-again, off-again relationship between mitochondrial genomes and species boundaries. Mol Ecol. 2017;26: 2212–2236. doi: 10.1111/mec.13959

65. Nelson TC, Muir CD, Stathos AM, Vanderpool DD, Anderson K, Angert AL, et al. Quantitative trait locus mapping reveals an independent genetic basis for joint divergence in leaf function, life-history, and floral traits between scarlet monkeyflower (*snp*) populations. bioRxiv. 2020;101: e02924–35. doi: 10.1101/2020.08.16.252916

66. Stankowski S, Streisfeld MA. Introgressive hybridization facilitates adaptive divergence in a recent radiation of monkeyflowers. Proc R Soc Lond B. 2015;282: 20151666. doi: 10.1098/rspb.2015.1666

67. Suarez-Gonzalez A, Hefer CA, Christe C, Corea O, Lexer C, Cronk QCB, et al. Genomic and functional approaches reveal a case of adaptive introgression from *Populus balsamifera* (balsam poplar) in *P. trichocarpa* (black cottonwood). Mol Ecol. 2016;25: 2427–2442. doi: 10.1111/mec.13539

68. Fontaine MC, Pease JB, Steele A, Waterhouse RM, Neafsey DE, Sharakhov IV, et al. Extensive introgression in a malaria vector species complex revealed by phylogenomics. Science. 2015;347: 1258524. doi: 10.1126/science.1258524

69. Figueiró HV, Li G, Trindade FJ, Assis J, Pais F, Fernandes G, et al. Genome-wide signatures of complex introgression and adaptive evolution in the big cats. Sci Adv. American Association for the Advancement of Science; 2017;3: e1700299. doi: 10.1126/sciadv.1700299

70. Angert AL, Schemske DW. The evolution of species’ distributions: reciprocal transplants across the elevation ranges of *Mimulus cardinalis* and *M. lewisii*. Evolution. 2005;59: 1671–1684. doi: 10.1554/05-107.1

71. Excoffier L, Foll M, Petit RJ. Genetic consequences of range expansions. Annu Rev Ecol Evol Syst. 2009;40: 481–501. doi: 10.1146/annurev.ecolsys.39.110707.173414

72. Wright S. Evolution in Mendelian populations. Genetics. 1931;16: 97–159.

73. Martin NH, Willis JH. Ecological divergence associated with mating system causes nearly complete reproductive isolation between sympatric *Mimulus* species. Evolution. 2007;61: 68–82. doi: 10.1111/j.1558-5646.2007.00006.x

74. Brandvain Y, Kenney AM, Flagel L, Coop G, Sweigart AL. Speciation and introgression between *Mimulus nasutus* and *Mimulus guttatus*. PLoS Genetics. 2014;10: e1004410. doi: 10.1371/journal.pgen.1004410

75. Hanson MR, Bentolila S. Interactions of mitochondrial and nuclear genes that affect male gametophyte development. Plant Cell. 2004;16 : S154–69. doi: 10.1105/tpc.015966

76. Charlesworth D, Ganders FR. The population genetics of gynodioecy with cytoplasmicgenic male-sterility. Heredity. 1979;43: 213–218.

77. Case AL, Finseth FR, Barr CM, Fishman L. Selfish evolution of cytonuclear hybrid incompatibility in *Mimulus*. Proc R Soc Lond B. 2016;283: 20161493. doi: 10.1098/rspb.2016.1493

78. Fishman L, Sweigart AL. When two rights make a wrong: the evolutionary genetics of plant hybrid incompatibilities. Annu Rev Plant Biol. 2018;69: 701–737. doi: 10.1146/annurev-arplant-042817-040113

79. Grossenbacher DL, Stanton ML. Pollinator-mediated competition influences selection for flower-color displacement in sympatric monkeyflowers. Am J Bot. 2014;101: 1915–1924. doi: 10.3732/ajb.1400204

80. Rastas P. Lep-MAP3: robust linkage mapping even for low-coverage whole genome sequencing data. Bioinformatics. 2017;33: 3726–3732. doi: 10.1093/bioinformatics/btx494

81. Catchen J, Amores A, Bassham S. Chromonomer: a tool set for repairing and enhancing assembled genomes through integration of genetic maps and conserved synteny. bioRxiv. 2020;32: 145–36. doi: 10.1101/2020.02.04.934711

82. Bolger AM, Lohse M, Usadel B. Trimmomatic: a flexible trimmer for Illumina sequence data. Bioinformatics. 2014;30: 2114–2120. doi: 10.1093/bioinformatics/btu170

83. Li H, Durbin R. Fast and accurate short read alignment with Burrows-Wheeler transform. Bioinformatics. 2009;25: 1754–1760. doi: 10.1093/bioinformatics/btp324

84. Li H, Handsaker B, Wysoker A, Fennell T, Ruan J, Homer N, et al. The Sequence Alignment/Map format and SAMtools. Bioinformatics. 2009;25: 2078–2079. doi: 10.1093/bioinformatics/btp352

85. McKenna A, Hanna M, Banks E, Sivachenko A, Cibulskis K, Kernytsky A, et al. The Genome Analysis Toolkit: a MapReduce framework for analyzing next-generation DNA sequencing data. Genome Res. 2010;20: 1297–1303. doi: 10.1101/gr.107524.110

86. Puzey JR, Willis JH, Kelly JK. Population structure and local selection yield high genomic variation in *Mimulus guttatus*. Mol Ecol. 2017;26: 519–535. doi: 10.1111/mec.13922

87. Altschul SF, Gish W, Miller W, Myers EW, Lipman DJ. Basic local alignment search tool. J Mol Biol. 1990;215: 403–410. doi: 10.1016/S0022-2836(05)80360-2

88. Quinlan AR, Hall IM. BEDTools: a flexible suite of utilities for comparing genomic features. Bioinformatics. 2010;26: 841–842. doi: 10.1093/bioinformatics/btq033

89. Li H. Tabix: fast retrieval of sequence features from generic TAB-delimited files. Bioinformatics. 2011;27: 718–719. doi: 10.1093/bioinformatics/btq671

90. Hoang DT, Chernomor O, Haeseler von A, Minh BQ, Vinh LS. UFBoot2: Improving the Ultrafast Bootstrap Approximation. Mol Biol Evol. 2018;35: 518–522. doi: 10.1093/molbev/msx281

91. Guindon S, Delsuc F, Dufayard J-F, Gascuel O. Estimating maximum likelihood phylogenies with PhyML. Methods Mol Biol. Totowa, NJ: Humana Press; 2009;537: 113–137. doi: 10.1007/978-1-59745-251-9_6

92. Stamatakis A. RAxML version 8: a tool for phylogenetic analysis and post-analysis of large phylogenies. Bioinformatics. 2014;30: 1312–1313. doi: 10.1093/bioinformatics/btu033

93. Mirarab S, Reaz R, Bayzid MS, Zimmermann T, Swenson MS, Warnow T. ASTRAL: genome-scale coalescent-based species tree estimation. Bioinformatics. 2014;30: i541–8. doi: 10.1093/bioinformatics/btu462

94. Green RE, Krause J, Briggs AW, Maricic T, Stenzel U, Kircher M, et al. A draft sequence of the Neandertal genome. Science. 2010;328: 710–722. doi: 10.1126/science.1188021

